# Identifying the causes and consequences of assembly gaps using a multiplatform genome assembly of a bird-of-paradise

**DOI:** 10.1101/2019.12.19.882399

**Authors:** Valentina Peona, Mozes P.K. Blom, Luohao Xu, Reto Burri, Shawn Sullivan, Ignas Bunikis, Ivan Liachko, Knud A. Jønsson, Qi Zhou, Martin Irestedt, Alexander Suh

**Affiliations:** Department of Ecology and Genetics – Evolutionary Biology, Uppsala University, Science for Life Laboratories, Norbyvägen 18D, SE-752 36, Uppsala, Sweden; Department of Organismal Biology – Systematic Biology, Uppsala University, Norbyvägen 18D, SE-752 36, Uppsala, Sweden; Department of Bioinformatics and Genetics, Swedish Museum of Natural History, SE-104 05, Stockholm, Sweden; Museum für Naturkunde, Leibniz Institut für Evolutions- und Biodiversitätsforschung, Berlin, Germany; MOE Laboratory of Biosystems Homeostasis & Protection, Life Sciences Institute, Zhejiang University, Hangzhou, China; Department of Molecular Evolution and Development, University of Vienna, Vienna, Austria; Department of Population Ecology, Institute of Ecology and Evolution, Friedrich-Schiller-University Jena, Dornburger Strasse 159, D-07743 Jena, Germany; Phase Genomics, Inc. 1617 8th Ave N, Seattle, WA 98109 USA; Uppsala Genome Center, Science for Life Laboratory, Dept. of Immunology, Genetics and Pathology, Uppsala University, SE-752 37, Uppsala, Sweden; Natural History Museum of Denmark, University of Copenhagen, Universitetsparken 15, DK-2100 Copenhagen, Denmark; Center for Reproductive Medicine, The 2nd Affiliated Hospital, School of Medicine, Zhejiang University

**Keywords:** Genome assembly, long reads, chromosome-level assembly, bird, transposable element, satellite repeat, GC content

## Abstract

Genome assemblies are currently being produced at an impressive rate by consortia and individual laboratories. The low costs and increasing efficiency of sequencing technologies have opened up a whole new world of genomic biodiversity. Although these technologies generate high-quality genome assemblies, there are still genomic regions difficult to assemble, like repetitive elements and GC-rich regions (genomic “dark matter”). In this study, we compare the efficiency of currently used sequencing technologies (short/linked/long reads and proximity ligation maps) and combinations thereof in assembling genomic dark matter starting from the same sample. By adopting different *de-novo* assembly strategies, we were able to compare each individual draft assembly to a curated multiplatform one and identify the nature of the previously missing dark matter with a particular focus on transposable elements, multi-copy MHC genes, and GC-rich regions. Thanks to this multiplatform approach, we demonstrate the feasibility of producing a high-quality chromosome-level assembly for a non-model organism (paradise crow) for which only suboptimal samples are available. Our approach was able to reconstruct complex chromosomes like the repeat-rich W sex chromosome and several GC-rich microchromosomes. Telomere-to-telomere assemblies are not a reality yet for most organisms, but by leveraging technology choice it is possible to minimize genome assembly gaps for downstream analysis. We provide a roadmap to tailor sequencing projects around the completeness of both the coding and non-coding parts of the genomes.

## Introduction

With the advent of Next Generation Sequencing (NGS) technologies, the field of genomics has grown exponentially and during the last 10 years the genomes of almost 10,000 species of prokaryotes and eukaryotes have been sequenced (from NCBI Assembly database, O’Leary et al. (2015)). Traditional NGS technologies rely on DNA amplification and generation of millions of short reads (few hundreds of bp long) that subsequently have to be assembled into contiguous sequences (contigs; Goodwin et al. (2016)). Although the technique has been revolutionary, the short-read length together with difficulties to sequence regions with extreme base composition poses serious limitations to genome assembly (Chaisson et al. 2015; Peona et al. 2018). Technological biases are therefore impeding the complete reconstruction of genomes and substantial regions are systematically missing from genome assemblies. These missing regions are often referred to as the genomic “dark matter” (Johnson et al. 2005). It is key now for the genomics field to overcome these limitations and investigate this dark matter.

Repetitive elements represent an important and prevalent part of the genomic dark matter of many genomes, given that their abundance and repetitive nature makes it difficult to fully and confidently assemble their sequences. This is particularly problematic when the read length is significantly shorter than the repetitive element, in which case it is impossible to anchor the reads to unique genomic regions. To what extent repeats can hamper genome assemblies depends on whether they are interspersed or arranged in tandem. Highly similar interspersed repeats, like for example transposable elements (TEs), may introduce ambiguity in the assembly process and cause assembly (contig) fragmentation. On the other hand, tandem repeats are repetitive sequences arranged head-to-tail or head-to-head such as microsatellites and some multi-copy genes (e.g., ribosomal DNA and genes of the Major Histocompatibility Complex, MHC). Reads shorter than the tandem repeat array will not resolve the exact number of the repeat unit, resulting in the collapse of the region into fewer copies. Some particular genomic regions enriched for repeats tend to be systematically missing or underrepresented in traditional genome assemblies. These regions include: 1) telomeres at the chromosome ends that are usually composed of microsatellites; 2) centromeres, essential for chromosome segregation often specified by satellites that can be arranged in higher-order structures like the alpha satellite in humans (Willard and Waye 1987) or by transposable elements in flies (Chang et al. 2019); 3) multi-copy genes like MHC genes (Shiina et al. 2009); d) non-recombining and highly heterochromatic chromosomes like the Y and W sex chromosomes (Chalopin et al. 2015; Smeds et al. 2015; Hobza et al. 2017). As these regions play an essential role in the functioning and evolution of genomes, the need to successfully assemble them is a pressing matter.

The other main limitation of traditional NGS methods is the shortcoming in reading regions with extreme base composition (an enrichment of either A+T or G+C nucleotides), thus representing another source of genomic dark matter. Extreme base composition mainly affects the last step of the standard library preparation for Illumina sequencers that involves PCR amplification (Dohm et al. 2008; Aird et al. 2011). GC-rich regions tend to have higher melting temperatures than the rest of the genome and are thus not as accessible with standard PCR protocols. On the other side of the spectrum, AT-rich regions are also challenging to be amplified with standard PCR conditions and polymerases (Oyola et al. 2012) because they require lower melting and extension temperatures (Su et al. 1996). Several protocols have been developed to help minimize the phenomenon of GC-skewed coverage (uneven representation of GC-rich regions), including PCR-free library preparation (Kozarewa et al. 2009) and isolation of the GC-rich genomic fraction prior to sequencing (Tilak et al. 2018). Nonetheless, there is no single method that entirely solves base composition biases of short-read sequencing and gives a homogeneous representation of the genome (Tilak et al. 2018). As a result, extremely GC-rich or AT-rich regions may not be assembled at all.

It is essential to be aware of technological biases and genome assembly incompleteness during project design since these can affect downstream analysis and mislead biological interpretations (Thomma et al. 2016; Weissensteiner et al. 2017; Domanska et al. 2018; Peona et al. 2018). For example, GC-skewed coverage is particularly important in birds, where 15% of genes are so GC-rich that they are often not represented in Illumina-based genome assemblies (Hron et al. 2015; Botero-Castro et al. 2017). Whether these genes are truly missing or mostly hiding due to technological limitations is still debated (Lovell et al. 2014, Botero-Castro 2017). However the “missing gene paradox” in birds is a clear example of how sequencing technologies can shape our view of genome evolution. Furthermore, some GC-rich sequences can form non-B DNA structures, i.e., alternative DNA conformations to the canonical double helix such as G-quadruplexes (G4). G4 structures are a four-stranded DNA/RNA topologies that seem to be involved into numerous cellular processes, such as regulation of gene expression (Du et al. 2008; Du et al. 2009; Raiber et al. 2011), genetic and epigenetic stability (Schiavone et al. 2014), and telomere maintenance (Biffi et al. 2012). On the repetitive element side, for example, transposable elements are a major target of epigenetic silencing (Law and Jacobsen 2010) that may influence the epigenetic regulation of nearby genes (Cowley and Oakey 2013; Chuong et al. 2016; Tanaka et al. 2019). The epigenetic effect of transposable elements may be beneficial or deleterious, but in either case it is important to acknowledge their potential involvement in the evolution of gene expression (Lerat et al. 2019). More generally, repetitive elements can play important roles in many molecular and cellular mechanisms, and as a source of genetic variability (Bourque et al. 2018). They have contributed to evolutionary novelty in many organismal groups, by giving rise to important evolutionary features like the mammalian placenta (Emera and Wagner 2012), the vertebrate adaptive immune system (Kapitonov and Koonin 2015; Zhang et al. 2019) and other telomere repair systems (Levis et al. 1993; McGurk et al. 2019). Thus, having genome assemblies that are as complete as possible facilitates research into a multitude of molecular phenomena (Slotkin 2018).

To achieve more complete genomes, we need new technologies. Recently, long-read single-molecule sequencing technologies with virtually no systematic error profile (Eid et al. 2009) have led to more complete and contiguous assemblies (English et al. 2012; Loomis et al. 2013; Pettersson et al. 2019; Smith et al. 2019). To date two sequencing strategies have been developed that produce very long reads from single-molecules: 1) Pacific Biosciences (PacBio) SMRT sequencing, in which the polymerases incorporate fluorescently labelled nucleotides and the luminous signals are captured in real time by a camera; 2) Oxford Nanopore Technologies, which sequences by recording the electrical changes caused by the passage of the different nucleotides through voltage sensitive synthetic pores. These new sequencing techniques have already yielded numerous highly contiguous *de-novo* assemblies (Faino et al. 2015; Gordon et al. 2016; Seo et al. 2016; Bickhart et al. 2017; Weissensteiner et al. 2017; Michael et al. 2018; Yoshimura et al. 2019) and helped improving the completeness of existing ones (Chaisson et al. 2014; Jain et al. 2018), as well as characterizing complex genomic regions like the human Y centromere and MHC gene clusters (Rhoads and Au 2015; Westbrook et al. 2015; Jain et al. 2018; Sedlazeck et al. 2018).

However, resolving entire chromosomes remains a difficult endeavour even with single-molecule sequencing (except for small fungal and bacterial genomes (Ribeiro et al. 2012; Thomma et al. 2016)). Even though no single technology is able to yield telomere-to-telomere assemblies, it is still possible to bridge separate contigs into scaffolds using long-range physical data and obtain chromosome-level assemblies. Scaffolding technologies are becoming more and more commonly used (Vertebrate Genome Project; Dudchenko et al. 2017; Belser et al. 2018; Deschamps et al. 2018; Li et al. 2019; Wallberg et al. 2019). The two most common ones are linked-reads (Weisenfeld et al. 2017) and proximity ligation techniques (reviewed in Sedlazeck et al. (2018)). Linked-read libraries are based on a system of labelling reads belonging to a single input DNA molecule with the same barcode (Weisenfeld et al. 2017). In this way, using high molecular weight DNA allows to connect different genomic portions (contigs) that may be distantly located but physically part of the same molecule. High-throughput proximity ligation techniques as Hi-C and CHiCAGO are able to span very distant DNA regions by sequencing the extremities of chromatin loops that could be up to Megabases apart in a linear fashion (for more details see Lieberman-Aiden et al. (2009)). While Hi-C is applied directly on intact nuclei, the CHiCAGO protocol reconstructs chromatin loops *in-vitro* from extracted DNA. All these libraries are then sequenced on an Illumina platform. As linked reads and proximity ligation techniques are becoming more and more popular used nowadays, we also implement and test them in the present study.

Although a plethora of new sequencing technologies and assembly methods are currently being successfully implemented, it remains unclear how they complement each other in the assembly process. Here we address these assembly and knowledge gaps using a bird as a model. Bird genomes represent a promising target to investigate that as their genomic features make it relatively easy to assemble most parts with the exception of few complex regions per chromosome. In fact, the typical avian genome is characterized by a small genome size (mean of ∼1 Gb Kapusta and Suh (2017); Gregory (2019)) and low overall repeat content (about 10% overall, with the exception of woodpeckers that have 20% (Kapusta and Suh 2017). However, there are gene-rich and GC-rich microchromosomes (Burt 2002; Griffin and Burt 2014; Miller and Taylor 2016) as well as a highly repetitive W chromosomes (at least in non-ratite birds Zhou et al. (2014); Smeds et al. (2015); Bellott et al. (2017)) that are still difficult to assemble.

In this study, to understand which genomic sequences are missing in regular draft genome assemblies with respect to a high-quality and curated assembly, we generated several draft *de-novo* genomes and a reference genome for the same sample of the paradise crow (*Lycocorax pyrrhopterus*, ‘lycPyr’). The paradise crow is a member of the birds-of-paradise family (Paradisaeidae), one of the most prominent examples of an extreme phenotypic radiation driven by strong sexual selection, and as such, a valuable system for the study of speciation, hybridization, phenotypic evolution and sexual selection (Shedlock et al. 2004; Irestedt et al. 2009; Ligon et al. 2018; Prost et al. 2019; Xu et al. 2019). We sequenced one female paradise crow individual with all the technologies that worked with a DNA sample of mean 50 kb molecule length. We combined short, linked, and long-read libraries together with Hi-C and CHiCAGO proximity ligation maps into a multiplatform reference assembly. All these technologies permitted us to curate the resulting assembly by controlling for consistency between multiple independent data types and make majority rule decision in conflicting cases. The curated assembly enabled us to: 1) demonstrate the feasibility of obtaining a high-quality assembly of a non-model organism with limited sample amount and non-optimal sample quality (a situation that empiricists commonly face); 2) identify which genomic regions are actually gained from combining technologies compared to draft assemblies of each individual technology; 3) assess the strengths and weaknesses of the implemented technologies regarding the efficiency of assembling difficult repeats and GC-rich regions; and 4) quantify how technologies can widen or limit the study of specific genomic features (e.g., TEs, satellite repeats, MHC genes, non-B DNA structures), thus providing a roadmap to investigate them.

## Results

We leveraged the power of data generated from multiple sequencing approaches for the same sample of paradise crow to generate a gold-quality assembly and to assess limitations of regular draft genomes based on any single technology. Briefly, we combined short, linked and long reads with proximity-ligation data to obtain a high-quality assembly despite the limitations of a non-model organism such as limited sample amount and non-optimal quality. For each sequencing technology, we produced an independent *de-novo* assembly. These assemblies were compared using majority-rule decisions by manually curating the final assembly. Finally, the multiplatform assembly was compared to each *de-novo* version to assess the amount of repeats and other complex regions previously missing from the individual assemblies. We then evaluated the completeness of each assembly using a variety of different metrics, including established scores such as BUSCO, contig/scaffold N50, LTR Assembly Index and new metrics like overall repeat content, number of MHC IIB exons, GC and G4 content, as well as number and nature of gaps.

### Long and short read *de-novo* assemblies

In order to compare the efficiency of short, linked, and long reads, we produced independent draft assemblies for each of the different sequence libraries. One draft genome assembly of *L. pyrrhopterus* based on short reads (Illumina) is already available from Prost et al. (2019) (‘lycPyrIL’; **Table 1**). For the present study, we produced two linked-read libraries (10X Genomics Chromium) from which we assembled two draft genomes (‘lycPyrSN1’ and lycPyrSN2’; where ‘SN’ stands for Supernova) and a PacBio library from the same paradise crow sample that generated the primary assembly ‘lycPyrPB’ (**Table 1** and **Methods** section). In total, four independent *de-novo* assemblies were generated.

**Table 1.** Draft and multiplatform assemblies generated for the paradise crow. For each assembly the sequencing technology and software used to produce them are shown together with contig N50, scaffold N50 and the number of gaps.

| Assembly | Technology | Software | Contig N50 (bp) | N contigs | Scaffold N50 (bp) | N scaffolds | N gaps<sup>a</sup> | Missing assembly<sup>b</sup> (%) |
| --- | --- | --- | --- | --- | --- | --- | --- | --- |
| **lycPyrIL** | Illumina HiSeq2500 (PE + MP)<sup>c</sup> | ALLPATHS-LG | 620,719 | 10,766 | 4,227,710 | 3,216 | 14,573 | 3.82 |
| **lycPyrPB** | PacBio RSII C6-P4 | Falcon | 6,644,420 | 3,422 | - | - | - | 0.45 |
| **lycPyrSN1** | 10X Genomics Chromium HiSeqX | Supernova2 | 144,856 | 29,791 | 4,360,585 | 13,934 | 21,550 | 4.53 |
| **lycPyrSN2** | 10X Genomics Chromium HiSeqX | Supernova2 | 149,640 | 27,366 | 4,748,626 | 14,217 | 20,131 | 2.62 |
| **lycPyrHiC** | PacBio + Phase Genomics Hi-C | Proximo | 6,644,420 | 3,422 | 70,588,898 | 2,927 | 533 | 0.45 |
| **lycPyrILPB** | lycPyrIL + gap-filling with PacBio | PBJelly | 1,982,606 | 6,895 | 4,229,628 | 3,216 | 10,422 | 3.03 |
| **lycPyr2** | PacBio + Dovetail CHiCAGO | HiRise | 6,294,665 | 3,463 | 6,644,037 | 3,227 | 282 | 0.45 |
| **lycPyr3** | lycPyr2 + 10X Genomics | ARCS + LINKS | 6,294,665 | 3,463 | 8,009,555 | 3,121 | 345 | 0.27 |
| **lycPyr4** | lycPyr3 + Phase Genomics Hi-C | Proximo | 6,294,665 | 3,463 | 69,071,023 | 1,713 | 1,791 | 0.27 |
| **lycPyr5** | lycPyr4 + manual curation with alignments + gap filling | PBJelly | 7,540,011 | 3,269 | 74,173,823 | 1,700 | 1,631 | 0.001 |
| **lycPyr6** | lycPyr5 + manual curation with Hi-C | Juicer | 7,540,011 | 3,271 | 74,173,823 | 1,700 | 1,635 | - |

<sup>a</sup> The number of gaps is estimated as the count of stretches of N nucleotides within a scaffold.
<sup>b</sup> The percentage of incompleteness is relative to the final version of the multiplatform assembly: $(1 - (\text{assembly size} / \text{final assembly size})) * 100$ . The N nucleotides are excluded from the calculation.
<sup>c</sup> PE: paired end reads; MR: mated reads.

We first evaluated the completeness of these assemblies by assessing their fragmentation, contig and scaffold N50 and by counting the number of core genes present with BUSCO (Nishimura et al. 2017; Waterhouse et al. 2017). In terms of fragmentation, the PacBio primary assembly (‘lycPyrPB’) consisted of about 3,000 contigs, while lycPyrIL had ∼3,000 scaffolds, and the 10XGenomics assemblies had about ∼14,000 scaffolds (**Table 1**). The short and linked-read assemblies all had a scaffold N50 of about 4 Mb while the PacBio assembly had a contig N50 of 6 Mb (**Table 1, Supplementary Table S1**). Notably, there is a 10-times higher of contig N50 in lycPyrPB relative to the lycPyrIL assembly, indicating significant improvement in assembly continuity in the PacBio vs. Illumina assembly. Next, we used the BUSCO tool (Nishimura et al. 2017) to identify correctly assembled core genes (percentage of only single-copy and complete genes follow): lycPyrIL 93.8%, lycPyrSN1 92.5%, lycPyrSN2 91.5%, lycPyrPB 84.8% prior to any assembly polishing (**Supplementary Table S2**). Similarly, we estimated genome completeness and quality of the intergenic and repetitive sequences with the LTR Assembly Index (LAI, Ou et al. (2018)). This index is calculated as the proportion of full-length LTR retrotransposons over the total length of full-length LTR retrotransposons plus their fragments. LAI could only be calculated for lycPyrPB since the other *de-novo* assemblies did not have enough complete LTR elements for the algorithm to work. lycPyrPB has an LAI score of 11.89, which is typical of a reference-quality assembly (Ou et al. 2018), and higher than chicken (galGal5, RefSeq accession number GCF_000002315.6; Bellott et al. (2017)) with an LAI score of 7.54. We cannot exclude that the higher score in paradise crow is caused by biological differences in LTR load between the species. More details about the LAI score distribution across chromosomes and genomes are found in **Supplementary Table S3, Supplementary Figure S1** and **Supplementary Figure S2**.

### The multiplatform reference assembly

To generate a high-quality genome assembly, we combined five technologies (short, linked, and long reads in addition to a CHiCAGO and Hi-C proximity ligation maps) into one multiplatform assembly. This process was divided into 9 steps (**Figure 1**), described in further detail in the **Methods** section.

**Figure 1.**
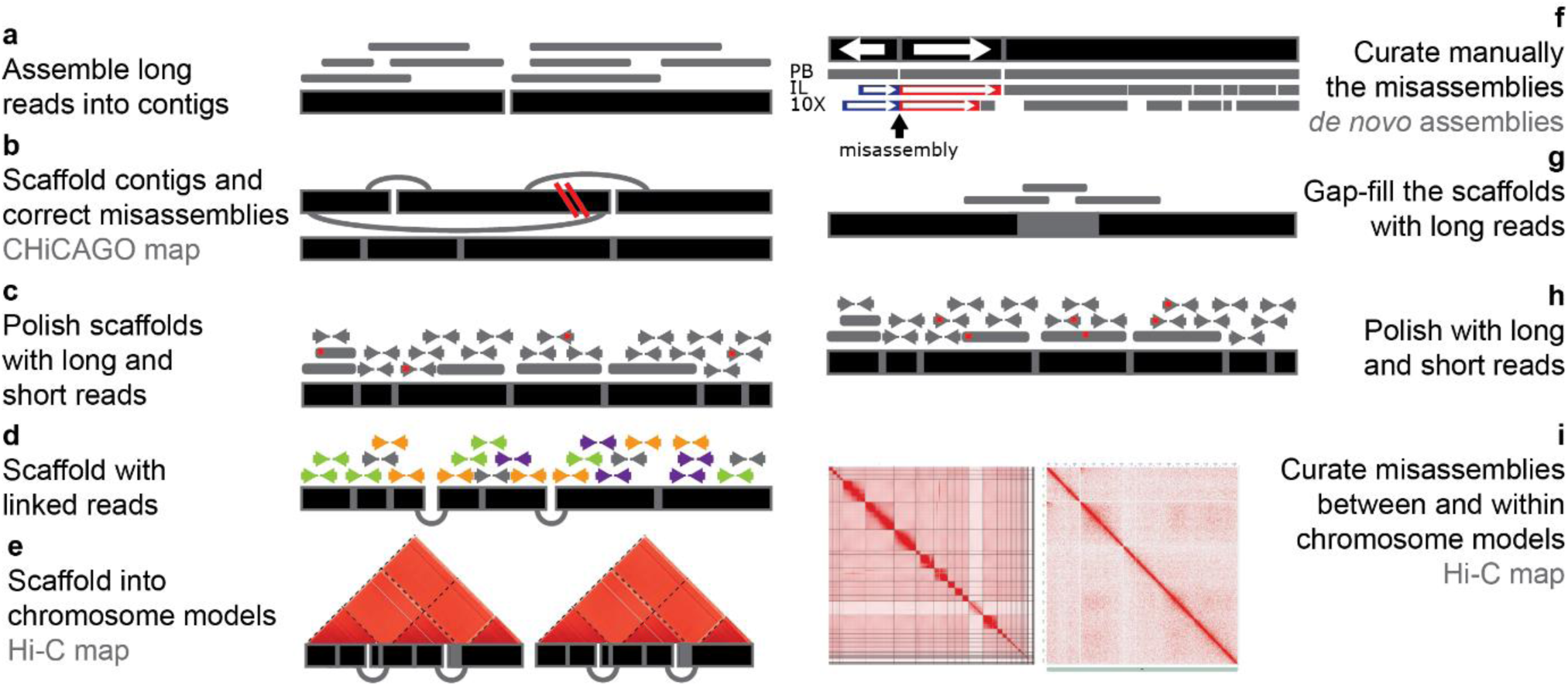
Overview of the multiplatform assembly process. **(a)** Long reads were assembled into contigs. (**b)** The primary assembly was corrected and scaffolded using long-range information provided by the CHiCAGO proximity ligation map. **(c)** The assembly was then polished from base-calling errors with both short and long reads and **(d)** further scaffolded with linked-reads. **(e)** The scaffolds are ordered and oriented into chromosome models according to the Hi-C proximity ligation map. **(f)** The chromosome models were aligned to the *de-novo* assemblies based only on one single technology and then manually inspected to find misassemblies and correct them following the majority rule (more details in **Figure 2** and **Methods**). PB: PacBio long-read assembly; IL: Illumina short-read assembly; 10X: 10XGenomics linked-read assemblies **(g)** Long reads were used to gap-fill the assembly and **(h)** to polish the final version together with short reads. **(i)** Hi-C heatmaps were used to identify and correct misassemblies between and within chromosome models.

First, we assembled the PacBio long reads into the primary assembly (lycPyrPB; 3,442 contigs) and it was scaffolded and corrected for misassemblies with the Dovetail CHiCAGO map (‘lycPyr2’; **Figure 1a-b**). The scaffolding software HiRise introduced 98 breaks and made 293 joins of scaffolds (gaps of 100 bp were introduced at this stage), as well as closed 11 gaps between contigs and resulted into an assembly of 3,227 scaffolds (**Table 1** and **Supplementary Table S1**). Subsequently we polished the assembly with long reads (two rounds of Arrow; Chin et al. (2016)) and short reads (two rounds of Pilon; Walker et al. (2014); **Figure 1c**).

We then continued to scaffold lycPyr2 with two types of long-range information in order to get a chromosome-level assembly. First, we used 10X Genomics linked reads (SN1 library; 24 kb mean molecule length; **Figure 1d**) that encode medium-range spatial information that placed 235 contigs into 131 new scaffolds. Of these new scaffolds we kept only 88 and discarded potential chimeric scaffolds, which were identified by being composed of sex-linked contigs and autosomal ones (based on male/female short-read coverage; see **Methods**). We then confirmed the chimeric nature of such scaffolds by constructing an additional assembly based on scaffolding lycPyrPB with the Hi-C map (‘lycPyrHiC’; **Table 1**). Phase Genomics Hi-C, i.e., 3D chromatin conformation data, can bridge sequences megabases apart (Burton et al. 2013) and theoretically reconstruct entire chromosomes (Hi-C super-scaffolds). In this way lycPyrHiC represented a second independent verification of the collinearity or chimeric nature of the contigs. Accordingly, we checked whether the contigs resided on different Hi-C super-scaffolds. Once we removed the chimeric contigs, we obtained ‘lycPyr3’ that contained a total of 3,121 scaffolds. Secondly, we scaffolded lycPyr3 with Phase Genomics Hi-C and obtained 38 super-scaffolds (‘lycPyr4’; **Figure 4e**) that harboured 1,446 contigs/scaffolds and accounted for 97% of the assembly, while 1,675 contigs/scaffolds remained unplaced (3%). As most of these super-scaffolds (32 out of 38) correspond to entire chromosomes of other avian species, we call them “chromosome models”. Examining the post-scaffolding Hi-C heatmap, we found that chromosomes 1 and 2 were split into two Hi-C super-scaffolds, respectively. Therefore, following the high level of Hi-C interaction between these super-scaffold pairs in the heatmap (**Supplementary Figure S3**), we manually combined the respective super-scaffold pair into one chromosome model (see **Methods**); the assembly thus resulted in 36 chromosome models.

**Figure 2.**
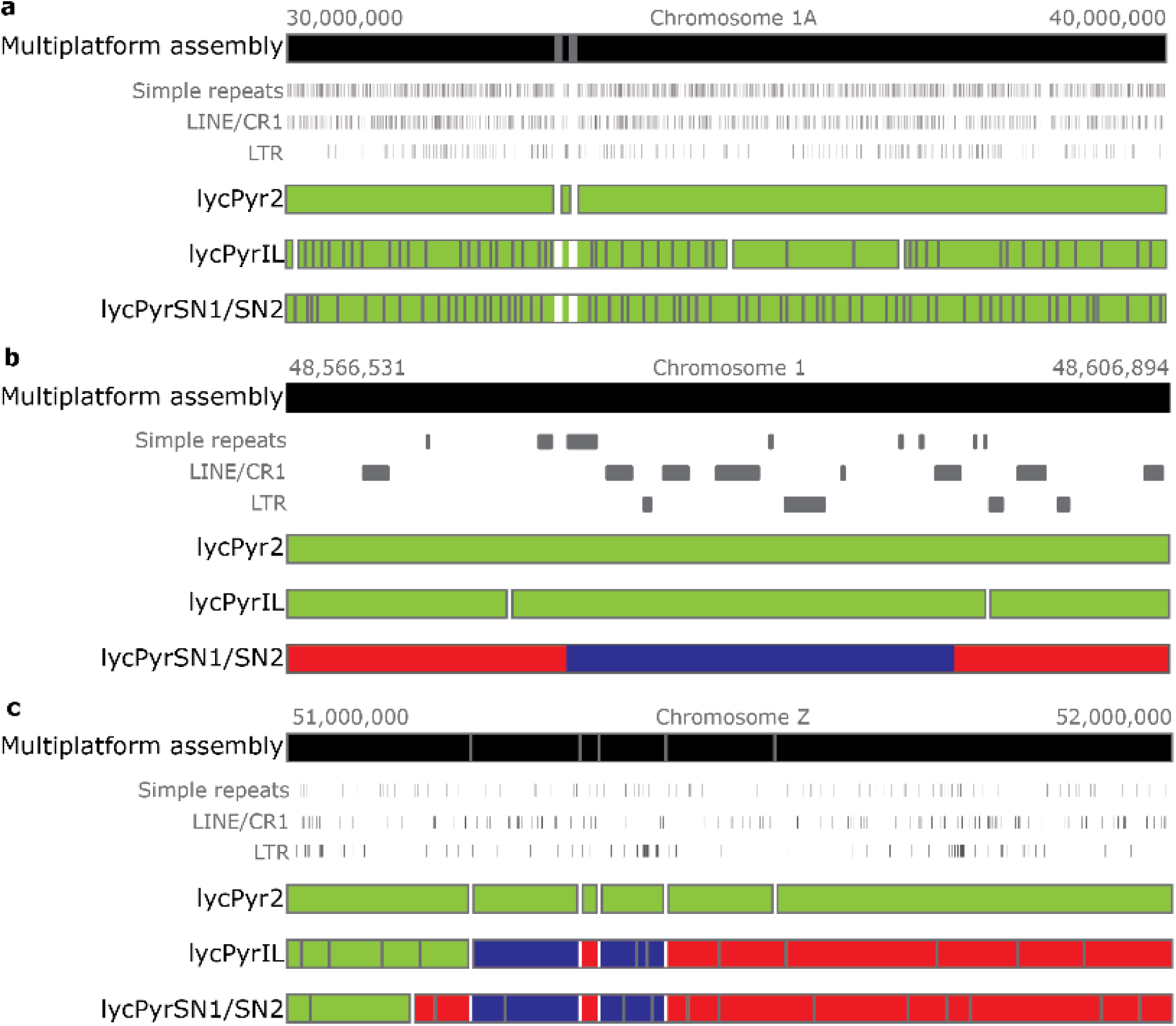
Examples of the manual curation of the assembly (**step f** in **Figure 1**). The multiplatform assembly is aligned to the other *de-novo* assemblies from the same sample. The grey lines within the assemblies represent gaps between different contigs or scaffolds while the white lines represent gaps within the same scaffold. Green means that the contigs/scaffolds align to the reference in the same orientation for their entire length while red and blue highlight contigs/scaffolds that partially align in the forward (red) and reverse (blue) direction to the reference. **(a)** Here 10 Mb of chromosome 1A are shown that are in accordance with all the *de-novo* assemblies. Nonetheless, short-read based technologies yielded much more fragmented scaffolds. **(b)** Example of a scaffold orientation misassembly in the 10XGenomics assembly. The other two assemblies span the inverted region and both agree with the multiplatform assembly. **(c)** Example of how two different assemblies could help to identify which contigs have to be re-oriented and re-ordered in the final assembly. In lycPyrIL, lycPyrSN1 and lycPyrSN2 we had scaffolds than span the misoriented (blue) region and bridge it to contigs that showed concordant orientation with the multiplatform assembly. This indicated that we have only a small local inversion of two PacBio contigs

**Figure 3.**
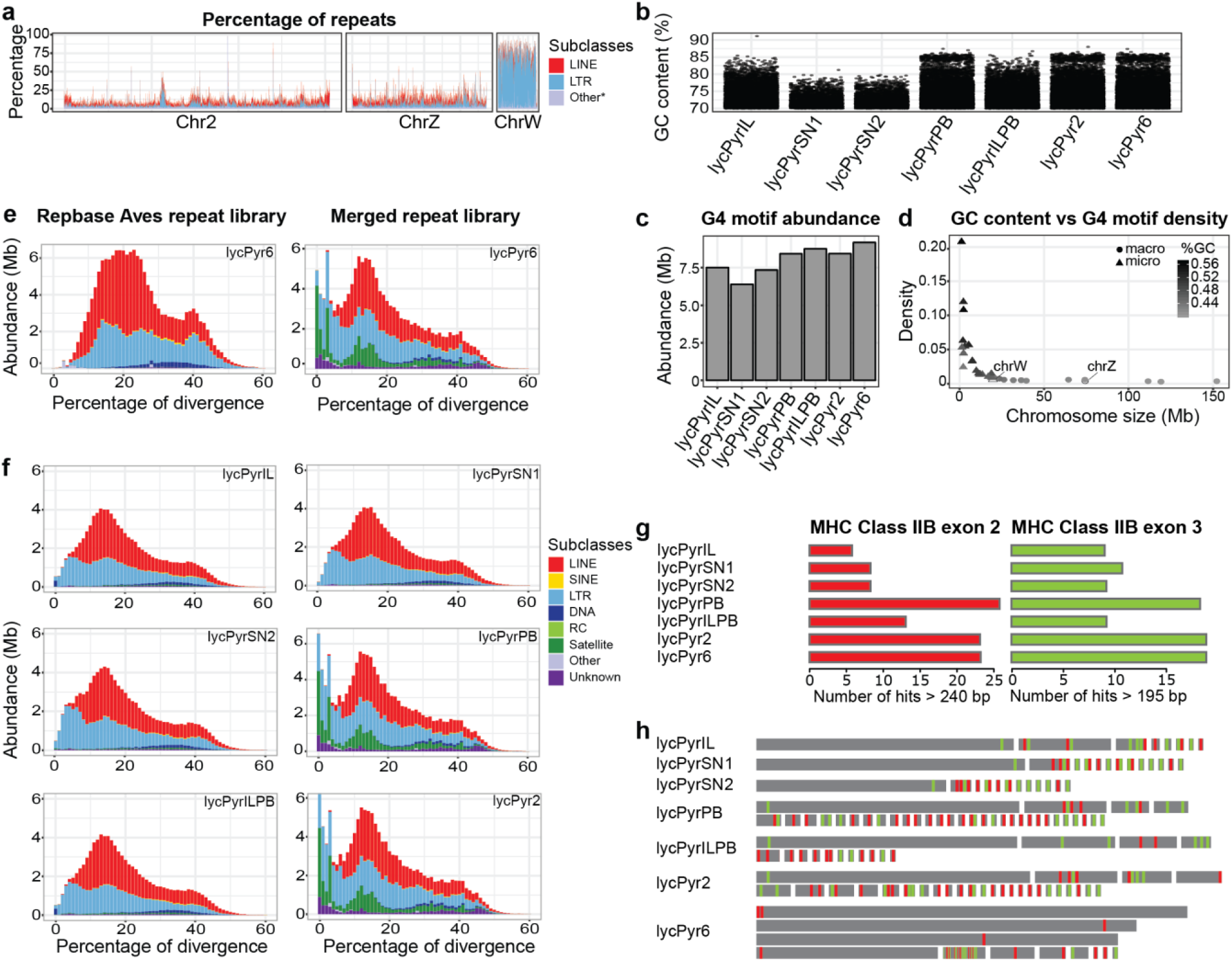
**(a)** Comparison of the repeat content across chromosome 2 (representative of autosomes), Z and W calculated as the percentage of repeats per window of 50 kb. Here LINE and LTR are shown as major components of the mobile element repertoire and all the other types of repeats are merged into the “Other*” category. **(b)** Distribution of GC-content per window (10 kb) across assemblies on the left side of the violin plots. GC-content distribution of the windows containing G4 motifs on the right side of the violin plots. **(c)** G4 motif abundance across different paradise crow assemblies. (**d**) G4 motif density across the chromosome models of the final assembly; the chromosomes are arranged by size; macrochromosomes are coloured in light grey while microchromosomes (smaller than 20 Mb) are shown in dark grey. The density distribution of G4 in micro and macro chromosomes was statistically different (t-test p-value: 0.01). **(e)** Repeat landscape of lycPyr6 masked with the Repbase Aves repeat library (on the left) and masked with the custom library produced in this study which also included the Repbase Aves library (on the right). **(f)** Repeat landscapes of the four *de-novo* assemblies of the paradise crow masked with the custom repeat library. **(g)** Abundance of MHC class IIB exon 2 and exon3 in the different paradise crow assemblies. **(h)** Schematic visualization of the instances of MHC class IIB exon 2 (red) and 3 (green). Each black rectangle represents a different contig or scaffold.

**Figure 4:**
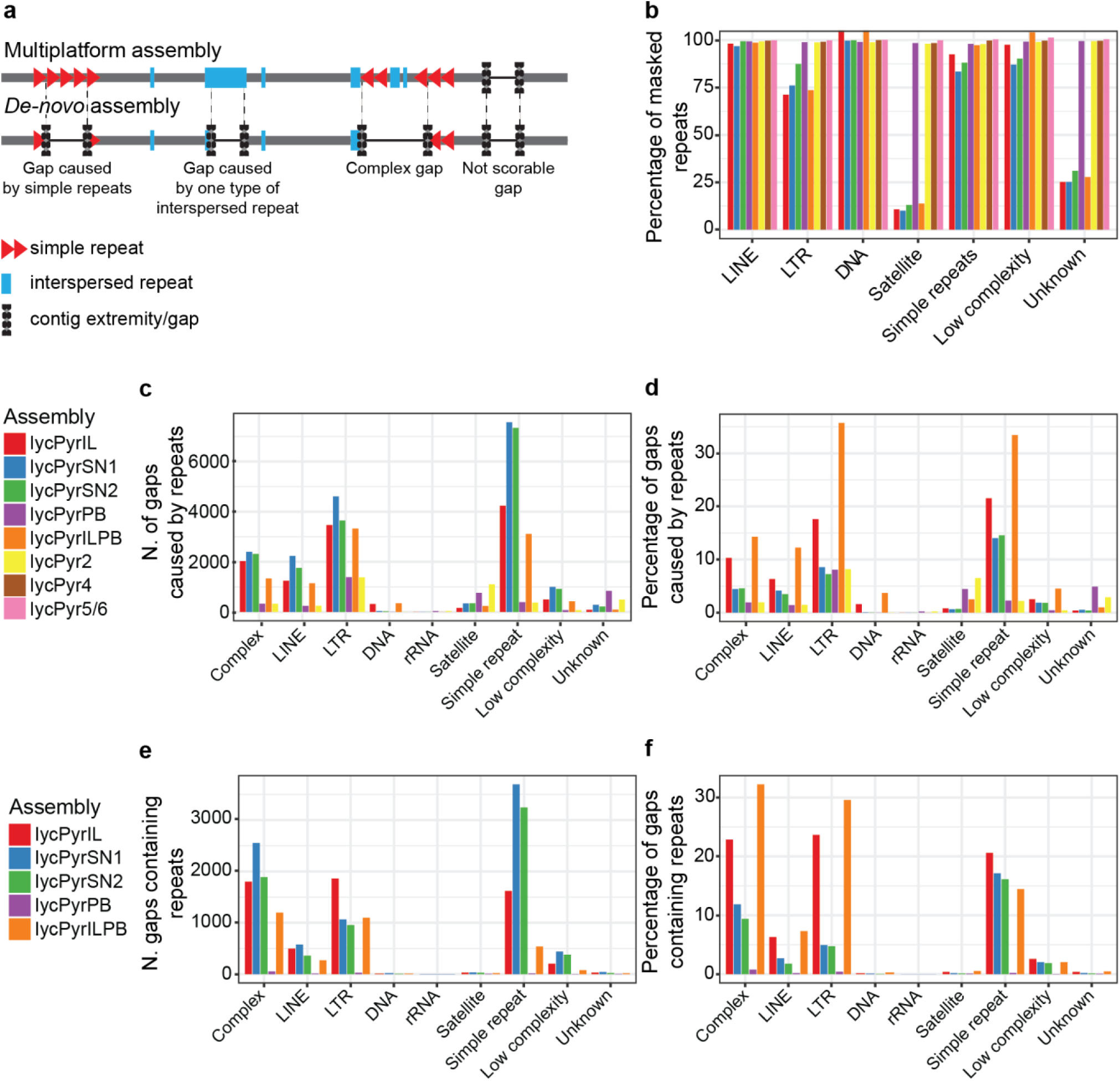
Overview of the causes and content of gaps in the paradise crow assemblies by comparing all the assembly versions to the final version. **(a)** Schematic representation of how gaps were categorized based on the flanking regions and content. **(b)** Proportion of repeats present in each assembly version respect to the reference (lycPyr6). **(c)** Number of gaps caused by the major repeat groups. **(d)** Proportion of gaps caused by the major repeat groups. (**e**) Number of gaps that contain (map to) repeats. (**f**) Proportion of gaps that contain (map to) repeats.

We proceeded to further manually curate the chromosome models by looking for misassemblies (**Figure 1f**) and used long reads for gap-filling (**Figure 1g**). We corrected fine scale orientation issues of contigs within scaffolds through whole genome alignments (see **Figure 2** and **Methods**) and corrected more orientation, order issues and erroneous chromosomal translocations through the inspection of Hi-C heatmaps (see **Figure 1i** and **Methods**). We first corrected 43 misassemblies by aligning the draft genomes and three outgroups to lycPyr4 (**Figure 2** and **Methods**). Next, we extended contig ends and filled scaffold gaps with long reads using PBJelly (‘lycPyr5’). PBJelly filled 106 gaps, extended 56 gaps on both ends and extended only one end of 292 gaps (**Supplementary Table S4**). Finally, we further checked for misassemblies with the help of the Hi-C data. We generated a Hi-C heatmap of lycPyr5 with Juicer (Durand et al. 2016) and detected misassemblies though the visual inspection of such a map with JuiceBox (Dudchenko et al. 2018) following the indications given by (Lajoie et al. 2015) and (Dudchenko et al. 2018). The Hi-C heatmap showed mostly orientation and ordering problems within lycPyr5 (**Supplementary Figure S4**) that can be identified from the ribbon-like patterns in the interaction map (Dudchenko et al. 2018). Finally, the map highlighted the misplacement of two contigs between chromosome models (**Supplementary Figure S4**). In total 76 misassemblies were corrected to generate the final assembly (‘lycPyr6’) with a scaffold N50 of ∼75 Mb (**Table 1**).

In parallel to the assembly of lycPyr6, we also generated a simpler multiplatform assembly by gap-filling the Illumina primary assembly (lycPyrIL) with PacBio reads (‘lycPyrILPB’). PBJelly was used to gap-fill the Illumina assembly and successfully closed 4,151 gaps, reducing the total number of gaps from 14,573 to 10,422. It also double extended 418 gaps and single extended 2,597 gaps (**Supplementary Table S4**). The numbers of scaffolds and scaffold N50 did not significantly change from lycPyrIL (**Table 1**).

### Chromosome models: macrochromosomes, microchromosomes and sex chromosomes

We obtained 36 chromosome models comprised of 16 macrochromosome models, 18 microchromosome models and two sex chromosome models. All the macrochromosome models showed homology to chicken chromosomes and were named after their homologous counterparts. The same applies for 12 of 18 microchromosomes, while the remaining 6 showed no homology with chicken chromosomes and therefore were tentatively named as unknown chromosomes “chrUN1-6”. The chromosomes homologous to chicken are mostly syntenic with respect to chicken with few exceptions. In fact, chicken chromosome 1 and 4 are split in two in Passeriformes and correspond, respectively, to chromosome 1 and 1A, and chromosome 4 and 4A (Kapusta and Suh 2017).

The Z and W sex chromosome models had an assembled size of 73.5 Mb and 21.4 Mb, respectively, and were comparable to chicken (82 Mb and 7 Mb, galGal6a, RefSeq accession number GCF_000002315.6; Bellott et al. (2017)). Z and W models were also largely consistent with the sex-linked contigs previously identified using male/female coverage comparisons (**Supplementary Table S5** and **Methods**), only 3.11 Mb of the W and 3.99 Mb of the Z chromosome were contigs not previously identified as sex-linked. Finally, the pseudoautosomal region (PAR) seemed to be fragmented into two parts. We identified two contigs that are homologous to the PAR of flycatcher; one of them was placed by Hi-C onto the Z while the other was placed onto the W chromosome model (**Supplementary Table S5**). While the Z chromosome showed a repetitive content similar to the autosomes (∼10%), the W was extremely repeat-rich (∼70%, **Figure 3a, Supplementary Table S6**). The dotplots of the alignments of the paradise crow sex chromosomes with the chicken sex chromosomes (**Supplementary Figure S5 and Supplementary Figure S6**) showed that the two Z chromosomes had a high level of synteny and collinearity while the repetitiveness of the two W chromosomes made it difficult to identify shared single-copy regions other than very small ones. The sex chromosomes were also easily identified in the post-clustering Hi-C heatmap (**Supplementary Figure S3**), as their hemizygosity can be expected to result in roughly half of the amount of Hi-C interactions (calculated as the frequency of shared paired-end reads between contigs/scaffolds) within each chromosome model and with the other chromosome models.

Finally, the LTR Assembly Index calculated on the single chromosomes yielded high scores (min 0 on chromosome 10, mean 13.14, max 21.41 on chromosome W) that have been suggested to be indicative of reference and gold-quality assemblies (Ou et al. (2018), **Supplementary Figure S1** and **Supplementary Table S3**).

### GC content and G4 motif prediction

GC-rich regions are commonly underrepresented in traditional NGS assemblies because of the aforementioned GC-skewed coverage phenomenon (see **Introduction**). Comparing the different *de-novo* assemblies, we noticed that indeed lycPyrPB showed more GC-rich regions (54,532 windows of 1 kb size with GC > 58.8%) with respect to lycPyrIL, SN1 and SN2 (45,966, 45,720 and 52,080 such windows, **Figure 3b, Supplementary Table S7, Supplementary Figure S7**). Thus, lycPyrSN1 shared a similar number of GC-rich regions while lycPyrSN2 was closer to lycPyrPB (**Supplementary Figure S7, Supplementary Table S7**).

Since GC-rich regions may form G-quadruplexes motifs and structures (G4), we expected the depletion of GC-rich short reads to limit the representation of G4 motifs in short read assemblies. Conversely, we expected G4 motifs to be more abundant in long read assemblies, since these have been suggested to be virtually free from sequence-based biases (Eid et al. 2009). To test this, we predicted the presence of G4 motifs using Quadron (Sahakyan et al. 2017) in all the different assemblies. All the *de-novo* Illumina-based assemblies had fewer predicted G4 sites the PacBio assemblies (**Figure 3c** and **Supplementary Table S8**). lycPyrSN2 and lycPyrIL had 7.3 and 7.5 Mb (169,214 and 166,602 motifs) occupied by G4 sequences and about 1.6 Mb or 24,000 motifs less than lycPyr6 (9.1 Mb, 193,248 motifs). lycPyrSN1 was the assembly with the fewest G4 motifs predicted (6.5 Mb, 149,275 motifs). The PacBio primary assembly lycPyrPB had 8.42 Mb of predicted G4, which was slightly higher in lycPyr2 after the correction with Dovetail CHiCAGO (8.43 Mb; **Figure 3c** and **Supplementary Table S8**). In the final assembly lycPyr6, G4 motifs were more present on microchromosomes than on macrochromosomes (**Figure 3d**).

### Repeat library

To obtain an in-depth annotation of interspersed and tandem repeats, the *de-novo* characterization of repetitive elements and manual curation thereof are essential (Platt et al. 2016). We manually curated a total of 183 consensus repeat sequences generated from lycPyrIL and lycPyrPB to have an optimal repeat characterisation. In Prost et al. (2019) a total of 112 raw consensus sequences were produced using RepeatModeler on three Illumina-based birds-of-paradise (*Astrapia rothschildii, L. pyrrhopterus* and *Ptiloris paradiseus*; including lycPyrIL) but only the 37 most abundant from lycPyrIL were manually curated. We then curated the remaining 75 and added 71 more *de-novo* consensus sequences based on curated raw consensus sequences from RepeatModeler run on lycPyrPB. Our new bird-of-paradise specific repeat library is now composed of the following numbers of consensus sequences: 56 ERVK, 56 ERVL, 37 ERV1, 5 CR1, 4 LTR, 9 satellites, 2 DNA transposons, 1 SINE/MIR, and 13 unknown repeats. All the consensus sequences curated for the three species of birds-of-paradise (*L. pyrrhopterus, A. rothschildii, P. paradiseus*) are given in **Supplementary Table S9**. Eventually, we merged birds-of-paradise consensus sequences together with the Repbase Aves library, the flycatcher (Suh et al. 2018), the blue-capped cordon blue (Boman et al. 2019) and the hooded crow libraries (Weissensteiner et al. 2019).

Custom and *de-novo* repeat libraries substantially improve the identification and masking of repeats in genome assemblies (Platt et al. 2016). To quantify this effect for our assemblies, we compared a general avian repeat library with our curated one. The custom library resulted in masking a higher fraction of the genome in every assembly (**Figure 3e-f**). When comparing the masked fraction with the custom library to the fraction masked with the Repbase library, we see that lycPyrIL, lycPyrILPB, and lycPyrSN1 have 20% more masked repeats (from 78 Mb to 94 Mb), while lycPyrSN2 has 21.68% (from 83 to 101 Mb), lycPyrPB 38% (from 87 Mb to 120 Mb), and lycPyr6 38% (from 88 Mb to 122 Mb; see **Figure 3e, Supplementary Table S10**). In particular, with the new library we were able to identify 9.4 Mb of satellite DNA in the PacBio-based assemblies, while the standard Repbase avian library identified only 1 Mb (**Figure 3e-f, Supplementary Table S10**). Relative to lycPyr6, most of the satellites and unknown repeats remain unassembled in the short-read and linked-read assemblies (**Figure 3f** and **Figure4b**).

### MHC class IIB analysis

In birds, the multi-copy gene family of the major histocompatibility complex (MHC) is arranged as a megabase long tandem repeat array (Miller and Taylor 2016). Since we expect it to be even more difficult to correctly assemble than the aforementioned interspersed repeats (O’Connor et al. 2019), it represents a prime candidate region for measuring the quality of an assembly.

We used the presence of entire copies of the second (most variable) and third (more conserved) exons of the MHC class IIB as proxies of assembly quality (Hughes and Yeager 1998). Overall, we found that short-read assemblies had fewer MHC gene copies than long-read assemblies (**Figure 3g-h**), while linked-read assemblies performed better than Illumina alone. Regarding exon 2 (**Figure 3g**), PacBio retrieved 26 copies while Illumina and 10XGenomics assembly only hold 6-8. However, it is worth noting that after correcting lycPyrPB with the Dovetail CHiCAGO map, 3 copies were lost (not detectable as full-length exons anymore) and were not restored by the subsequent steps of sequence corrections and curation. The results were similar for exon 3 (**Figure 3g**): PacBio assemblies retrieved 18-19 copies while the other technologies retrieved only 9-11 copies. In this case we see that the molecule input length of 10XGenomics library has an effect on the assembly of these genes, where the library with shorter molecule length had assembled more copies than the longer one (11 vs 9 exon 2 copies; **Figure3g-h**). On the other hand, while Dovetail CHiCAGO prevented the identification of some exon 2 copies, it increased the number of assembled copies of exon 3.

### Gap analysis

The process of scaffolding links contigs together without adding any information about the missing DNA between them, but it is possible to use long reads to fill those gaps. For this we utilized PBJelly (English et al. 2012) to extend and bridge contigs in the assembly by locally assembling PacBio reads to the contig extremities. Once the software finds reads aligned to the contig extremities, the extremities can be: 1) extended on one or both sides to reduce the gap length, 2) extended and bridged to fill the entire gap, 3) extended over the length of the gap without being ultimately bridged (overfilled). PBJelly extended the extremities of 348 gaps, closed 116 gaps and overfilled 236 gaps (**Supplementary Table S4**). This gap-filling step added a total of 2.96 Mb to the assembly. All the sequences that were extended or gap-filled were more GC-rich (40%-89%, mean 58%) than the average GC content of 40% and 2865 G4 motifs were added for a total of 171 kb. Only 800 kb of the 2.96 Mb added were repetitive elements; specifically, ∼400 kb of LTR elements were added, 120 kb of LINE, 142 kb of satellite DNA and 90 kb of simple and low complexity repeats (**Supplementary Table S4**).

Furthermore, we investigated the causes of assembly fragmentation in several assemblies by analysing the immediate adjacency of repetitive elements to the gaps (lower part of **Figure 4a**). We found that simple repeats were the major fragmentation cause in Illumina and 10XGenomics assemblies, followed by LTR and LINE elements (**Figure 4c-d**). In contrast, PacBio gaps (lycPyrPB and lycPyr2) seemed to be mainly caused by LTR elements and secondarily by satellites (**Figure 4c-d**).

Finally, we quantitatively and qualitatively assessed which repeats in the final multiplatform assembly lycPyr6 were collapsed as gaps in the draft assemblies (**Figure 4e-f**). Many gaps in the Illumina and 10XGenomics draft assemblies corresponded to complex regions consisting of multiple types of repetitive elements (**Figure 4e-f**). Among draft assembly gaps containing only a single type of repeat in lycPyr6, most were caused by simple repeats, LTR retrotransposons, and LINE retrotransposons in short-read and linked-read assemblies (**Figure 4e-f**).

## Discussion

Assembling complete eukaryotic genomes is a complex and demanding endeavour often limited by technological biases and assembly algorithms (Alkan et al. 2010; Sedlazeck et al. 2018). In the last decade, NGS technologies defined the standard of genome assemblies. Although they provided an unprecedented view on the structure and evolution of many coding regions (Zhang et al. 2014), short reads hardly inform on the entire complexity of a genome (Thomma et al. 2016). Indeed, the systematic absence from genome assemblies and the difficulty to characterize the nature of many such genomic regions (e.g. centromeres, telomeres, other repeats and highly heterochromatic regions) gave these “unassemblable” sequences the evocative name of genomic “dark matter” (Johnson et al. 2005; Weissensteiner and Suh 2019).

In this study, we demonstrated that a combined effort involving multiple state-of-the-art methods for long-read sequencing and scaffolding yielded a high-quality reference for a non-model organism. We showed that a multiplatform approach was highly successful in resolving elevated quantities of genomic dark matter in respect to single-technology assemblies (regular draft assemblies) and thus resulted in a much more complete assembly. In order to assess genome completeness we focused mostly on the quantification and characterization of previously inaccessible regions within genomic dark matter, such as large transposable elements, GC-rich regions, and the high-copy MHC locus.

We generated a *de-novo* multiplatform assembly of a female bird-of-paradise genome by combining the cutting-edge technologies that are now being implemented in many assembly projects (Faino et al. 2015; Gordon et al. 2016; Seo et al. 2016; Bickhart et al. 2017; Weissensteiner et al. 2017; Michael et al. 2018; Yoshimura et al. 2019), namely Illumina short reads, 10XGenomics linked reads, PacBio long reads and two proximity ligation maps with Dovetail CHiCAGO and Phase Genomics Hi-C. The choice of using a bird-of-paradise is manifold. First, avian genomes are small among amniotes and have an overall repeat content of 10%, which make most genomic regions relatively “easy” to assemble. This has made it possible to focus on regions that are challenging to assemble in eukaryotic genomes of any size and complexity, like the repeat-rich W sex chromosome, and the GC-rich microchromosomes. Second, birds-of-paradise is a highly promising system for the study of speciation, hybridization and sexual selection (Irestedt et al. 2009; Prost et al. 2019; Xu et al. 2019). A gold standard genome for this family will consequently expose new possibilities for more in-depth studies of the genomic evolution behind the spectacular radiation of birds-of-paradise.

By employing a multiplatform approach, we 1) could assemble a chromosome-level genome which includes the W chromosome and several previously inaccessible microchromosomes (i.e., comparable to the chicken genome, so far the best avian genome available); 2) report that a substantial proportion (up to 90%) of repeat categories like satellites and LTR retrotransposons are missing from most types of *de-novo* assemblies (**Figure 3e-f, Figure 4b**); and 3) identify simple repeats and LTR retrotransposons as the major causes of assembly fragmentation (**Figure 4c-d**).

### A chromosome-level assembly for a non-model organism

Our final assembly comprises 36 chromosome models. This assembled chromosome number is similar to the known karyotype of another bird-of-paradise species *Ptiloris intercedens* (36-38 chromosome pairs; Les Christidis, personal communication). Among these models, there are 16 macrochromosomes, 12 microchromosomes, and the Z and W sex chromosomes showing homology to chicken chromosomes (galGal6a). The remaining 6 models do not share homology with known chicken chromosomes (galGal6a) and they might be putatively uncharacterized microchromosomes. Microchromosomes are known to be very GC-rich (Burt 2002) and indeed this trend is present in our data as well (**Figure 3d**). Base composition can create biases during the sequencing process especially when a PCR step is required for the library preparation (Dohm et al. 2008; Aird et al. 2011) thus limiting the representation of GC-rich and AT-rich reads in the data. Although, long read sequencing technologies like PacBio have reduced amplification-based biases to a minimum (Schadt et al. (2010) but see Guiblet et al. (2018)), we could not assemble contiguous sequences for all microchromosomes. Among the unknowns and unassembled chromosomes, chromosome 16 which is one of the most complex avian chromosomes and also holds the MHC (Miller and Taylor 2016). The absence of these chromosomes is likely explained by that they are by far the densest in G4 motifs of all chromosomes (**Figure 3d**). Given that DNA polymerase tends to introduce sequencing errors in the presence of G4 structures (Guiblet et al. 2018), it is tempting to think that the depletion of microchromosomes from assemblies is not only due to GC content per se but also due to the potential presence of non-B structures (like G4) that elevated GC content appears to correlate with. Nonetheless, even with the extensive use of cytogenetics the last chicken assembly (galGal5; Warren et al. (2017)) completely lacks 5 microchromosomes. It thus seems plausible that these chromosomes need special efforts to be recovered.

One of the most surprising outcomes of this multiplatform approach is the successful assembly of the highly repetitive W chromosome which turned out to be larger (assembly size 21 Mb) and more repetitive than the chicken equivalent (assembly size 9 Mb; Bellott et al. (2017)). In both species, it is likely that the assembled sequences cover the euchromatic portions of the W. Birds have a ZW sex chromosome system where the female is the heterogametic sex and the female-specific W is analogous to the mammalian male-specific Y chromosome. Comparable to the mammalian Y (Charlesworth et al. 2000), the W chromosome is highly repetitive and difficult to assemble (Weissensteiner and Suh 2019). Previous studies focusing on the repetitive content of the avian W in chicken (Bellott et al. 2017) and collared flycatcher (Smeds et al. 2015) showed in both cases a repeat density of about 50%. In our assembly of the paradise crow, we found the W chromosome to be even more repetitive with a repeat density of ∼70% and highly enriched for LTR retrotransposons (**Figure 3a** and **Supplementary Table S6**). Having assembled chromosomes is key to improve any genomic analysis but studies on sex chromosome evolution in birds has so far been heavily biased towards Z (Zhou et al. 2014; Yazdi and Ellegren 2018; Xu et al. 2019). With genome assemblies like the present, it will be possible to improve reconstructions how the two sex chromosomes diverged. We can already see that the W chromosome evolves rapidly (**Supplementary Figure S5**) via accumulation of transposable elements and only few regions appear syntenic between paradise crow and chicken W.

### How complete are genome assemblies?

Previous studies (see for example Etherington et al. (2019); Paajanen et al. (2019)) have assessed the efficiency of available sequencing technologies in genome assembly and genome completeness mainly through summary statistics like scaffold N50 and BUSCO. Scaffold N50 indicates the minimum scaffold size among the largest scaffolds making up half of the assembly, while BUSCO values measure the number of complete/incomplete/missing core genes in the assembly. However, genome completeness goes beyond scaffold N50 and gene presence (Thomma et al. 2016; Domanska et al. 2018; Sedlazeck et al. 2018). Genes usually occupy a small fraction of genomes and new sequencing technologies commonly yield high N50 values. Therefore, these statistics have a very limited scope in perspective of what the new sequencing technologies can achieve.

Although often being used as proxy of assembly quality, scaffold N50 is hardly meaningful in this regard since it does not inform about the completeness and correctness of the assembled sequences. If we order the scaffolds by decreasing size, scaffold N50 value can only reflect the fragmentation level of the first half of the assembly regardless of whether the second half is made up of shorter sequences. Finally, contig N50 should be used as a measure of contiguity, rather than scaffold N50, as contig length measures sequences not interrupted by gaps.

Most of the currently available avian genomes score more than 94% of BUSCO gene completeness (Peñalba et al. 2019) with various degrees of fragmentation, suggesting that it has become straightforward to generate short-read assemblies with high BUSCO values. On the other hand, BUSCO seems to be limited by the sequencing errors introduced by PacBio in the identification of gene models (Watson and Warr 2019). Even with multiple rounds of error correction, BUSCO fails to recognize genes that are actually present, at least partially, in the assembly (Watson and Warr 2019). Moreover, BUSCO seems to be trained and based on a set of core genes identified from Sanger and Illumina assemblies. As such, BUSCO does not quantify genes in PacBio assemblies that were previously missing in Illumina genomes, which would be needed for a fair genome completeness comparison. This tendency is also evident from our results: for example after gap-filling lycPyrIL with long reads, 10 genes were not detectable anymore in the resulting assembly lycPyrILPB (**Supplementary Table S2**). A similar dynamic was observed also during the assembly process of the superb fairy-wren *Malarus cyaneus* (Peñalba et al. 2019) where BUSCO values dropped with long-read gap-filling but were restored after sequence polishing.

The new technologies have the potential to assemble very repetitive regions (e.g. MHC) and elusive chromosomes (e.g., W and microchromosomes). For this reason, quality assessment should rely upon measuring the efficiency in assembling difficult regions and not on those regions that we already obtain with previous technologies. We therefore decided to measure genome completeness and quality by characterising and quantifying repetitive regions.

Long reads were instrumental, not only to find and mask more repeats, but also to assemble and discover previously overlooked repetitive sequences. In fact, by adding PacBio sequence data we were able to significantly increase the number of predicted repeat subfamilies compared to the repeat library previously built on three birds-of-paradise species (from 112 to 183 consensus sequences; Prost et al. (2019)). These 71 new consensus sequences were only predicted by RepeatModeler using the PacBio assembly, probably because the respective repeats were too fragmented or assembled in too few copies in Illumina assemblies. A clear example is given by the satellite DNA repeats that are severely depleted from both the lycPyrIL assembly (**Figure 3e-f, Figure4b**) and from the previous repeat library. With our new repeat library we could increase the base pairs masked by RepeatMasker by up to 38 % within the same assembly (lycPyr6). This indicates that while longer read lengths are important for assembling repeats, only with a comprehensive repeat library we can quantify their actual efficiency.

Repetitive elements are not only made up of transposable elements and satellite repeats, but also of multi-copy genes. One of the most repetitive gene family is the Major Histocompatibility Complex (MHC) involved in the adaptive immune response. In birds, MHC genes are located on one of the most difficult chromosomes to assemble, namely chromosome 16 (Miller and Taylor 2016). We recovered several scaffolds from this chromosome for which the only, though fragmented, assembly exists from chicken (Warren et al. 2017). We counted how many MHC IIB copies we could retrieve in the different assemblies, using BLAST hits to exon 2 and 3 sequences as proxy. We found the maximum number of copies in lycPyrPB (**Figure 3g-h**) followed by lycPyr6, suggesting that the misassembly correction with the CHiCAGO map affected the MHC genes, with the number of hits of exon 2 decreasing and for exon 3 increasing. Short-read assemblies harbour fewer MHC IIB exon copies but we note that 10XGenomics could assemble a couple more copies compared to standard Illumina data. Moreover, lycPyrSN1 contained slightly more MHC genes than lycPyrSN2 assembled with longer input molecule length.

As a further use of repetitive elements as quality measures, we tested the LTR Assembly Index (LAI; Ou et al. (2018)) that assesses the quality of an assembly from the completeness of the LTR retrotransposons present. It was not possible to obtain values for the Illumina and 10XGenomics assemblies because the tool requires a certain baseline quantity of the full-length LTR assembled to run as initial requirements. Nonetheless, both lycPyrPB and lycPyr6 show LAI scores (respectively 11.89 and 13.59, **Supplementary Table S3, Supplementary Figure S1**) typical for high-quality reference genomes (as indicated in Ou et al. (2018)) and higher than those of chicken (**Supplementary Figure S2**). The increase in LAI value from lycPyrPB and lycPyr6 indicates that the assembly curation process, mostly gap-filling and polishing, improved the quality of the primary assembly.

In addition to repetitive elements, base composition is the other main factor that limits completing genome assemblies. We thus assessed the GC-content per window for each assembly (**Figure 3b, Supplementary Figure S7**) and as expected, found more GC-rich windows in lycPyrPB compared to the other *de-novo* assemblies (**Supplementary Figure S7**). High GC-content is often associated with non-B DNA structures like G4 that have been shown to introduce sequencing errors during polymerisation (Guiblet et al. 2018). We predicted the presence of G4 motifs in our assemblies (**Figure 3c**) and Illumina and 10XGenomics assemblies have about 1.6-2.6 Mb less of G4 compared to lycPyrPB. In this case, linked reads did not help to get a more complete overview of this genomic feature respect to regular Illumina libraries. On the other hand, the overall curation from lycPyrPB to lycPyr6 improved G4 prediction. G4 structures influence various molecular mechanisms such as alternative splicing and recombination, therefore more complete assemblies make these regions accessible for comparative genomic analysis.

### Strengths and limitations of sequencing technologies

Nowadays, we have a plethora of sequencing technologies to choose from, each with their own advantages and limitations. On top of that, the large number of assembly tools available and hundreds of parameters to tweak makes it inevitable to produce numerous different assembly versions. For example, we generated 15 different assemblies only for the parameter optimization of the linked-read scaffolding (**Figure 1d**) and there are studies generating even 400 assemblies in total (Montoliu-Nerin et al. 2019). In such a situation, it might seem difficult to decide how to choose the “best” assembly among dozens. Here we present what we learned from the different technologies and how they help in resolving the genomic regions that are most difficult to assemble.

We used two types of *de-novo* assemblies based on Illumina sequencing. The first, lycPyrIL is an Illumina assembly made from multiple insert size libraries of paired end and mate pair reads (Prost et al. 2019); the second on 10XGenomics linked reads (lycPyrSN1 and SN2). It is notable that lycPyrIL is much more contiguous than lycPyrSN1 and SN2 (contig N50 of 620 kb vs 145-150 kb; **Table 1**) and has much fewer gaps. Although lycPyrIL is a less fragmented assembly, lycPyrSN2 has a better resolution for repeats since 7 Mb more repeats are masked and a larger number of MHC IIB exons are present (**Figure3g-h**) as well as G4 motifs (**Figure 3g**). Nonetheless, the contiguity reached in lycPyrPB for the same sample at contig level (contig N50 of 6 Mb) is ten-fold higher than in lycPyrIL and even outscores lycPyrIL scaffold N50 of 4 Mb. 10XGenomics linked reads bring long-range information through the barcode system that is useful for local phasing, detection of structural variations (Zheng et al. 2016; Marks et al. 2019), scaffolding (Yeo et al. 2017) and construction of recombination maps (Dréau et al. 2019; Sun et al. 2019). We used the barcode information to scaffold the PacBio assembly (lycPyr3, **Table 1**) without obtaining many new scaffolds but this could be due to the already high contiguity of the input lycPyrPB assembly. Finally, we note that the molecule input length for the 10XGenomics libraries have different effects on the assembly and BUSCO scores. That is, lycPyrSN1 (24 kb mean molecule length library) outscores lycPyrSN2 (26.1 kb mean molecule length library) in the number of complete BUSCO genes (**Supplementary Table S2**). Even though 10XGenomics linked reads consist of short reads, both lycPyrSN1 and lycPyrSN2 have more missing genes compared to lycPyrIL (**Supplementary Table S2**).

Long reads together with proximity ligation maps are game changers in genomics. Their combination yielded a very high-quality assembly for a non-model bird with suboptimal sample quality (see mean molecule lengths for 10XGenomics assemblies above). The PacBio assembly is by far the most contiguous and a suitable genomic backbone to obtain chromosome models including the W chromosome and several microchromosomes. The main weakness linked to PacBio is the introduction of sequencing errors (mostly short indels) that must be corrected with accurate short reads. As mentioned before, the sequencing errors hinder the identification of gene models (BUSCO) and protein prediction (Watson and Warr 2019). Moreover, the PacBio assembly is likely not free of misassemblies (e.g., chimeric contigs). Thus a second type of independent data is necessary to detect such errors; e.g., ∼100 potential misassemblies were identified by the CHiCAGO proximity map. The CHiCAGO map was very useful to correct the assembly and make a first scaffolding, but neither alone nor with 10XGenomics scaffolding yielded a chromosome-level assembly. The only type of data implemented here that allowed the generation of chromosome models was the Hi-C map. The latter does not rely on extracted DNA quality or library insert size, but instead on *in-situ* proximity within the nuclei of the fixed sample. As such, Hi-C data is an effective replacement of linkage maps for scaffolding purposes (Dudchenko et al. 2017) and can be used to manually curate assemblies.

A direct way to identify the limits of sequencing data is to investigate where assemblers fail to resolve sequences, i.e. where contig fragmentation occurs. Therefore, we characterized what causes contig fragmentation in each assembly by analysing sequences directly adjacent to gaps and inferring the gap content of draft assemblies by aligning their flanks to the final multiplatform version lycPyr6 (**Figure 4a**). In general, we found that long and/or homogeneous repeats such as LTR retrotransposons, satellites, and simple repeats are the main fragmentation causes in every assembly, though the specific repeat type changed with the technology. Short-read and linked-read contigs mostly break at simple repeats. Even though the percentage of simple repeats assembled in lycPyrIL, lycPyrSN1 and lycPyrSN2 ranges between 80-90% relative to lycPyr6 (**Figure 4b**), simple repeats also caused most of the assembly gaps, indicating that insert size and linked read methods are not sufficient to unambiguously solve those regions (**Figure 4c-d**). At the same time, the gaps of these three assemblies, when compared to the final multiplatform assembly, mainly contain LTR retrotransposons, simple repeats and complex repeats (defined as arrays of different types of repeats; **Figure 4e-f**). LTR retrotransposons are the second most abundant retrotransposons in the paradise crow assembly and several kilobases long. These features make LTR retrotransposons the major cause of fragmentation in the PacBio assembly and the second in the short-read ones. This partially unexpected trend is likely because LTR retrotransposons are underrepresented in lycPyrIL, lycPyr SN1 and lycPyrSN2 (as indicated by their lack of part of the recent LTR activity; **Figure 3e-f**). The same pattern can be observed for the multicopy rRNA genes: the only assemblies showing gaps caused by rRNA genes are the PacBio-based and this is likely because PacBio was the only technology able to (partially) solve those repeats (**Figure 4c-d**). It is interesting that linked reads appear to better distinguish long repeats like LTR retrotransposons than short-read libraries based on insert size (**Figure 4b**). The satellite portion of the genome was significantly better assembled with PacBio long reads (∼9 Mb), while neither multiple Illumina libraries nor linked reads could assemble more than 1 Mb of satellites. This is probably due to the highly homogeneous nature of long stretches of satellites that make satellite arrays collapse during assembly (Hartley and O’Neill 2019). Similar to LTR retrotransposons and rRNA genes, satellites are barely assembled in lycPyrIL, lycPyrSN1 and lycPyrSN2. Therefore satellites are not a major cause of contig fragmentation in Illumina-based assemblies. LINEs are usually short retrotransposons due to 5’ truncation during integration (Levin and Moran 2011)and in the paradise crow and other songbirds they seem to be mostly present in old copies (**Figure 3e**; Suh et al. (2018); Weissensteiner et al. (2019)). Therefore they likely are less homogeneous elements, with more diagnostic mutations and hence easier to assemble. In fact, both Illumina and 10XGenomics assemblies have 96-98% of LINEs assembled and LINEs represent only the fourth causative factor of fragmentation. Finally, we noticed a disproportion of DNA transposons annotated in the Illumina assemblies (lycPyrIL and lycPyrILPB) compared to the other assemblies. This phenomenon might be explained by annotation issues linked to the fragmentation of those regions or by the presence of unsolved haplotypes. DNA transposons have been inactive in songbirds for even longer than LINEs (Kapusta and Suh 2017) and should thus be rather straightforward to assemble.

## Conclusions

Thanks to a manually curated multiplatform assembly and three *de-novo* draft assemblies for the same sample, we were able to characterise and measure genome completeness across sequencing technologies. As expected, long-read assemblies are more complete than short-read assemblies but completeness has been usually measured with statistics that are optimized for short reads rather than for long reads. Scaffold N50 and BUSCO values do not reflect the entire potential and strengths of new sequence technologies, therefore we measured completeness focusing on the most difficult-to-assemble genomic regions. By doing so, we traced the essential steps for generating a high-quality assembly for a non-model organism while optimizing costs and efforts.

Based on our assembly comparisons, the essential elements to make a chromosome-level assembly are a contiguous primary assembly based on long reads, an independent set of data for correcting misassemblies (CHiCAGO map or linked reads) and polish sequencing errors (short or linked reads), and a Hi-C map for chromosome-level scaffolding. PacBio needs error correction both at the nucleotide level (base calling errors and short indels) and at the assembly level (e.g., chimeric contigs). For both scopes it is possible to use Illumina data but a note of caution is due. First, when polishing the assembly for base calling errors and short indels, short reads could over-homogenize repetitive sequences and thus it would be advisable to correct only outside repeats. In addition, 10XGenomics linked reads can also be used to correct both sequencing errors and misassemblies (e.g., Tigmint, Jackman et al. (2018)) and to scaffold the genome (ARCS, Yeo et al. (2017), ARKS, Coombe et al. (2018), fragScaff, Adey et al. (2014)). In general, the spatial information brought by linked reads seems to be very versatile (e.g., assembly correction, scaffolding, structural variation inference, haplotype phasing) and able to better avoid over-collapsing of repetitive elements and genes (**Figure 3** and **4**). Therefore, if budgets and sample material are limited, this technology may be suitable to obtain a better genomic overview than short reads alone. Nevertheless, long reads provide the most detailed look into difficult-to-assemble genomic regions. We summarized the strengths and limitations of the implemented technologies in **Figure 5** that can be used as a guide for choosing technologies and ranking assemblies.

**Figure 5.**
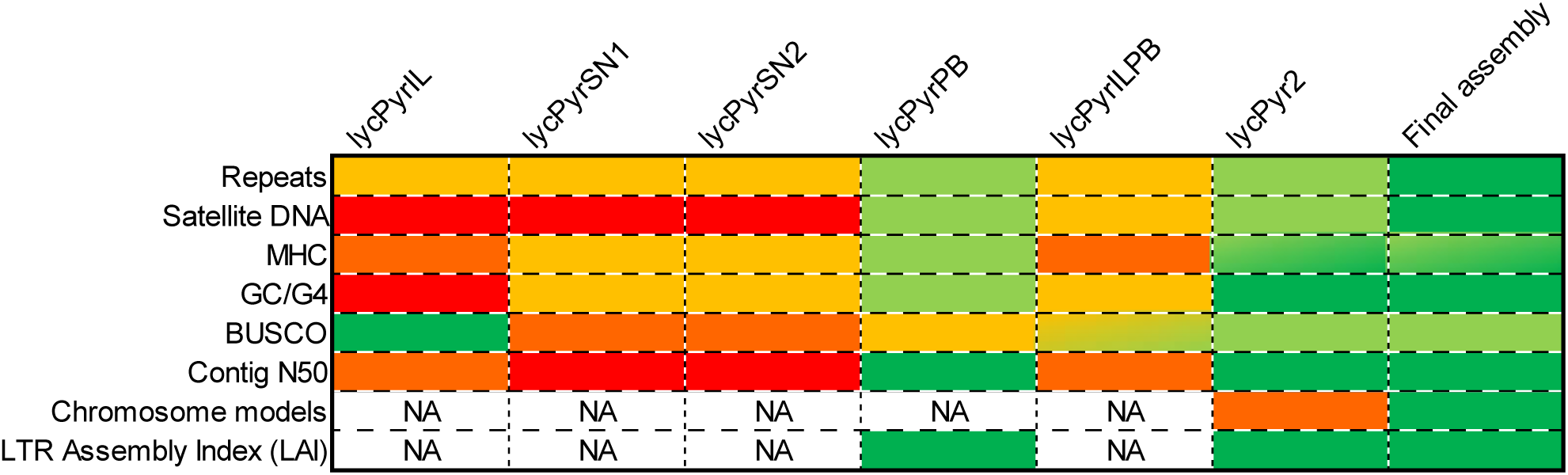
Summary of the relative efficiency of the different technologies over quality/completeness parameters. Green: most effective; red: least effective.

We have shown that recent technological developments have led to enormous improvements in assembly quality and completeness, paving the way to more complete comparative genomic analyses, including regions that were previously inaccessible within genomic dark matter. At the same time, awareness of technological strengths and weaknesses in resolving repeat-rich and GC-rich regions is fundamental for choosing the most suitable technology when designing sequencing projects, and will help in a dilemma many genome scientists face these days: choosing the best assembly among many.

## Methods

### Samples

We used pectoral muscle samples from three vouchered specimens of *Lycocorax pyrrhopterus* ssp. *obiensis* collected on Obi Island (Moluccas, Indonesia) in 2013, from the Museum Zoologicum Bogoriense (MZB) in Bogor, Indonesia, temporarily on loan at the Natural History Museum of Denmark. One female (voucher: MZB 34.073) sample preserved in DMSO was used for PacBio, Illumina and 10XGenomics sequencing and for the Dovetail CHiCAGO library, one female sample (voucher: MZB 34.070) preserved in RNAlater was used for the Hi-C library with Phase Genomics, and one male sample preserved in DMSO (voucher: MZB 34.075) was used for Illumina sequencing.

### Sequencing technologies and *de-novo* assemblies

We sequenced the female sample MZB 34.073 using a) PacBio RSII C6-P4 (mean of 11 kb and N50 of 16 kb for read length) for a total coverage of 72X; b) 10XGenomics with a HiSeqX Illumina machine (24 kb mean molecule length, 280 bp library insert size, 150 bp read length, net coverage 39.7X); c) 10XGenomics with HiSeqX Illumina machine (26.1 kb mean molecule length, 280 bp library insert size, 150 bp read length, net coverage 37.9X). DNA was extracted using magnetic beads on a Kingfisher robot, except for library c) which was based on DNA extracted with agarose gel plugs as in (Weissensteiner et al. 2017). In addition to these libraries, we also used the Illumina libraries and assembly produced in (Prost et al. 2019): Illumina HiSeq 2500 TruSeq paired-end libraries (180 bp and 550 bp insert sizes) and Nextera mate pair libraries (5 kb and 8 kb insert sizes) for a total coverage of 90X. Furthermore, two paired-end libraries (125 bp read length) of chromatin-chromatin interactions from CHiCAGO and Hi-C techniques were produced using a HiSeq 2500 by Dovetail Genomics (Putnam et al. 2016) and Phase Genomics (more details below), respectively. Finally, we generated a paired-end library with insert size of 650 bp on an Illumina HiSeqX machine for the male sample.

For each library/technology (namely Illumina, 10XGenomics and PacBio) we made independent *de-novo* assemblies. (Prost et al. 2019) used ALLPATHS-LG (Butler et al. 2008) for Illumina data while we used Falcon (Chin et al. 2016) for PacBio data and Supernova2 (Weisenfeld et al. 2017) for 10XGenomics data (**Table 1**). All the basic genome statistics of the assemblies (**Supplementary Table S1**) were calculated using the Perl script assemblathon_stats.pl from https://github.com/KorfLab/Assemblathon/blob/master/assemblathon_stats.pl.

### Identification of sex-linked contigs and PAR

Given the extreme conservation of the Z chromosomes of songbird (Xu et al. 2019), we used the Z-chromosome sequence of great tit as a query to search for homologous Z-linked contigs in paradise crow. The aligner nucmer was used to perform the one-to-one alignment of the great tit genome and lycPyrPB. Those contigs with more than 60 percent sequence aligned to great tit Z chromosome were identified as Z-linked. We further calculated the sequencing coverage using the female Illumina paired-end libraries to confirm the half-coverage pattern of candidate Z-linked contigs relative to autosomal contigs. We used BWA-MEM to map the reads and the samtools depth function to estimate contig coverage. To identify candidate W-linked contigs, we calculated the re-sequencing coverage of the male individual, because W-linked contigs are female-specific and are not expected to be mapped by male reads while the coverage of female reads should be half of that of autosomes. We used the known PAR sequences of collared flycatcher (Smeds et al. 2014) to identify the homologous PAR contigs in paradise crow. As expected, the PAR contigs were found to show similar re-sequencing coverage in both the male and the female as on the autosomes.

### Multiplatform approach

We created three types of multiplatform assemblies, one that combines only Illumina and PacBio data (lycPyrILPB, see **Table 1**), a second one combining PacBio and Hi-C data, and a third more comprehensive one that combines three types of sequencing data and two types of proximity ligation data (lycPyr6).

For the first type of assembly (lycPyrILPB), we used the Illumina assembly lycPyrIL (Prost et al. 2019) as genomic backbone and gap-filled it with PacBio long reads using the software PBJelly (PBSuite v. 15.8.24) maintaining the all the default options but -min 10 to consider only gaps of at least 10 base pairs length. The second multiplatform assembly lycPyrHiC was built by scaffolding the PacBio primary assembly (lycPyrPB) with Hi-C data.

For the most comprehensive assembly (lycPyr6), we combined PacBio, Illumina, 10XGenomics, CHiCAGO and Hi-C data (**Figure 1**). The first step was to assemble the PacBio reads into a primary assembly with the Falcon software (Chin et al. (2016); **Figure 1a**). The primary contigs were corrected and scaffolded with the Dovetail CHiCAGO map generating lycPyr2 (**Figure 1b**) using the software HiRise (Putnam et al. 2016). lycPyr2 then was polished with long reads (two runs of Arrow; Chin et al. (2016)) and short reads (three runs of Pilon 1.22; Walker et al. (2014); **Figure 1c**). Since PacBio sequencing is prone to introduce short indels in the reads (Eid et al. 2009), we addressed specifically these sequencing problems with Pilon while we did not correct single nucleotide variants. Furthermore, in order to not over-polish repetitive regions (i.e., homogenising them with short reads), we excluded Pilon corrections falling within repeats identified by RepeatMasker 4.0.7 using our custom repeat library.

We then scaffolded lycPyr2 using the long-range information given by 10XGenomics linked reads with the software ARCS 1.0.1 (Yeo et al. (2017); parameters -s 95 -e 1000 -m 20-100000) and LINKS 1.8.5 (Warren et al. (2015); parameters -a 0.2) generating lycPyr3 (**Figure 1d**). The parameters for ARCS and LINKS have been chosen after generating 15 assemblies with different values for -m -e -a (**Supplementary Table S11**). The optimal parameter combination was established by minimising a) the number of “private” scaffolds belonging only to one combination of parameters, b) the number of scaffolds containing putative in-silico chromosomal translocations.

lycPyr3 was scaffolded into chromosome models (clusters of contigs and scaffolds) with the Phase Genomics Hi-C data and the Proximo Hi-C scaffolding pipeline (lycPyr4; **Figure 1e**). Hi-C data were generated using a Phase Genomics (Seattle, WA) Proximo Hi-C Animal Kit. Following the manufacturer’s instructions for the kit, intact cells from two samples were crosslinked using a formaldehyde solution, digested using the Sau3AI restriction enzyme, and proximity ligated with biotinylated nucleotides to create chimeric molecules composed of fragments from different regions of the genome that were physically proximal *in-vivo*, but not necessarily linearly proximal. Continuing with the manufacturer’s protocol, molecules were pulled down with streptavidin beads and processed into an Illumina-compatible sequencing library.

Reads were aligned to the draft assembly lycPyr3 following the manufacturer’s recommendations. Briefly, reads were aligned using BWA-MEM (Li and Durbin (2010); v. 0.7.15-r1144-dirty) with the -5SP and -t 8 options specified, while keeping the other parameters as default. SAMBLASTER (Faust and Hall 2014) was used to flag PCR duplicates, which were later excluded from analysis. Alignments were then filtered with samtools (Li et al. (2009); v1.5, with htslib 1.5) using the -F 2304 filtering flag to remove non-primary and secondary alignments, as well as read pairs in which one or more mates were unmapped. Phase Genomics’ Proximo Hi-C genome scaffolding platform was used to create chromosome-scale scaffolds from the draft assembly as described in (Bickhart et al. 2017). As in the LACHESIS method (Burton et al. 2013), this process computes a contact frequency matrix from the aligned Hi-C read pairs, normalised by the number of Sau3AI restriction sites (GATC) on each contig, and constructs scaffolds in such a way as to optimise expected contact frequency and other statistical patterns in Hi-C data. Approximately 286,000 separate Proximo runs were performed to optimise the number of scaffolds and scaffold construction in order to make the scaffolds as concordant with the observed Hi-C data as possible.

Two chromosomes (chr1 and chr2) appeared to be split into two different super-scaffolds (or clusters) respectively, thus they were manually put together following the orientation suggested by the Hi-C interaction heatmap (**Supplementary Figure S3**). We then manually inspected the assembly lycPyr4 for misassemblies (**Figure 1f** and **Figure 2**) by aligning the four *de-novo* assemblies (lycPyrIL, lycPyrPB, lycPyrSN1 and lycPyrSN2) to it using Satsuma2 (Grabherr et al. 2010) and chromosome models from three songbird outgroups (*Ficedula albicollis, Taeniopygia guttata* and *Parus major*) with LASTZ 1.04.00 (Harris 2007). We identified misassemblies by looking for regions in which the different *de-novo* assemblies were in conflict with the final assembly (schematically showed in **Figure 2**). We applied the majority rule for each scaffolding or orientation conflict found between lycPyr4 and the four draft assemblies. To make any decisions against the scaffold configuration in lycPyr4, three of the four *de-novo* assemblies needed to be in discordance with lycPyr4 and show the same pattern of discordance. In cases where only two *de-novo* assemblies showed the same pattern of discordance and the other were not informative, we used the information provided by the outgroups to decide whether to keep the lycPyr4 scaffold configuration or correct it. With this approach we were able to identify 45 intra-scaffold misassemblies at a fine scale, all of them being orientation issues of PacBio contigs within scaffolds.

Then, we gap-filled the assembly using PBJelly (PBSuite 15.8.24; English et al. (2012)) with the default options except for the parameter -min 10 in order to consider the gaps longer than 10 bps (**Figure 1g**). After the gap-filling step that used the PacBio reads, we ultimately polished the genome with long reads using Arrow (one run; PacBio library) and with short reads using Pilon (two runs; Illumina library; **Figure 1h**).

The last step of assembly curation involved the generation of Hi-C heatmaps on lycPyr5 by mapping the Hi-C library to the assembly using Juicer 1.5 (Durand et al. (2016); **Figure 1i**). We manually inspected the Hi-C maps for misassemblies using Juicebox 1.9.8 (https://github.com/aidenlab/Juicebox) and corrected lycPyr5 accordingly (**Supplementary Figure S3**). This way, we manually solved remaining assembly issues regarding the orientation and order of some contigs or scaffolds within the chromosome models, as well as corrected *in-silico* chromosomal translocations.

The completeness of the assemblies was assessed with gVolante (Nishimura et al. 2017) using BUSCO v3 for avian genomes (**Supplementary Table S2**) and with LTR Assembly Index (Ou et al. (2018); **Supplementary Table S3**).

The mitochondrial genome was identified as one PacBio contig by aligning the mtDNA of *Corvus corax* (GenBank accession number KX245138.1) to lycPyrPB. It was annotated using DOGMA (Wyman et al. 2004) and tRNAscan-SE 1.3.1 (Lowe and Eddy (1997); **Supplementary Table S12**).

### Chromosome nomenclature

Since the chicken genome is the best avian genome assembled so far with reliable chromosome information (Warren et al. 2017), we named and oriented our chromosome models according to homology with galGal5 (RefSeq accession number GCF_000002315.6). In the case that our chromosome models were not completely collinear with chicken, we oriented them following the orientation of the majority of the model respect to chicken. Finally, if the chromosome models did not share any homology with chicken, their orientation was not changed.

### Repeat library

We produced a *de-novo* repeat library for paradise crow by running the RepeatMasker 4.0.7 and RepeatModeler 1.0.8 software on the PacBio *de-novo* assembly. We hard-masked lycPyrPB with the Aves repeat library from Repbase (version 20170127; https://www.girinst.org/about/repbase.html) together with the consensus sequences from (Prost et al. 2019), then ran RepeatModeler. The new consensus sequences generated by RepeatModeler were aligned back to the reference genome; the 20 best BLASTN 2.7.1+ results were collected, extended by 2 kb on both sides and aligned to one another with MAFFT 7.4.07. The alignments were manually curated applying the majority rule and the superfamily of repeat assessed following the (Wicker et al. 2007) classification.

All the new consensus sequences were masked in CENSOR (http://www.girinst.org/censor/index.php) and named according to homology to known repeats in the Repbase database. Sequences with high similarity to known repeats for their entire lengths were given the name of the known repeat + suffix “_lycPyr”; repeats with partial homology have been named with the suffix “-L_lycPyr” where “L” stands for “like” (Suh et al. 2018). Repeats with no homology with known ones have been considered as new families and named with the prefix “lycPyr” followed by the name of their superfamilies.

The final repeat library also contains the manually curated version of the consensus sequences previously generated on other two birds-of-paradise *Astrapia rothschildi* “astRot”, *Ptiloris paradiseus* “ptiPar” (Prost et al. 2019), the ones from *Corvus cornix* (Weissensteiner et al. 2019), *Uraeginthus cyanocephalus* (Boman et al. 2019), *Ficedula albicollis* and all the avian repeats available on Repbase (mostly from chicken and zebra finch).

### G4 motif identification

The *de-novo* assemblies and the final version have been scanned for G-quadruplex (G4) motifs with the software Quadron (Sahakyan et al. (2017); https://github.com/aleksahak/Quadron). Only non-overlapping hits with a score greater than 19 were used for subsequent analysis as suggested in (Sahakyan et al. 2017). The density of such motifs per chromosome model was calculated using bedtools coverage (BEDTools 2.27.1; Quinlan (2014)).

### MHC class IIB analysis

To infer how highly duplicated genes are assembled with different input data and assembly strategies, we investigated the distribution of major histocompatibility class IIB (MHCIIB) sequence hits in seven assemblies: lycPyrIL, lycPyrPB, lycPyrSN1, lycPyrSN1, lycPyrILPB, lycPyr2 and lycPyr6 (the intermediate assemblies like lycPyr3 are not shown here because the MHC content did not change from lycPyr2 to lycPyr5). We performed BLAST (Altschul et al. 1990) searches both with sequences of the highly variable exon 2 that encodes the peptide binding region, and with the much more conserved exon 3 (Hughes and Yeager 1998), as the disparate levels of polymorphism within these regions may provide insights into different aspects of challenges with genome assembly. We conducted tBLASTn (BLAST 2.7.1+) searches using alignments available from Goebel et al. (2017) that include sequences from across the entire avian phylogeny. We chose this strategy to ensure the identification MHCIIB sequences, as with sequences of only a single-species BLAST search might miss highly divergent sequences as they are often present in the MHC, where within-species diversity of MHC genes often equals between-species divergence. From the available alignments, we exclusively retained sequences spanning the entire 270 bp of exon 2 and sequences covering 220 bp of exon 3. This left query alignments including 233 sequences from 22 bird orders/families for exon 2, and 314 sequences from 26 bird orders/families for exon 3. Overlapping blast hit intervals were merged. To ensure that these intervals contained sequences corresponding to MHCIIB, we first BLAST searched them back against the GenBank database using BLASTn queries, and retained only intervals producing hits with MHCIIB. We then aligned the remaining sequences using the MAFFT alignment server with the --add option and default settings, and manually screened the alignments to identify non-MHCIIB sequences. Finally, we determined the alignment lengths of BLAST hit intervals after removing insertions relative to the query alignment. We report only hits longer than 240 bp for exon 2 and longer than 195 bp for exon 3, corresponding to approximately 90% of the respective query alignment lengths.

### Gap analysis

For each assembly produced, we estimated the number of gaps caused by repeats by intersecting the gap and repeat coordinates using bedtools window (Quinlan 2014) with a window size of 100 bp (**Figure 4a**). Only gaps longer than 10 bp were taken into consideration. This filter is particularly important for lycPyrIL since there are many small gaps of 1-5 Ns that are probably caused by sequencing or base-calling errors.

We estimated what is missing in the draft assemblies with respect to the final multiplatform assembly lycPyr6 by aligning the flanking regions to the gaps onto the final version. We then assessed the presence of annotated repeats on lycPyr6 between the aligned flanking regions to the draft assembly gaps. To do these pairwise alignments, we extracted 500 bp of flanking regions from the intra-scaffold gaps of lycPyrIL, lycPyrSN1, lycPyrSN2, lycPyrPB and lycPyrILPB and BLASTn searched the sequences to lycPyr6 with BLAST 2.7.1+. The alignments were filtered to retain only unambiguously orthologous positions on lycPyr6, namely there was only one alignment (98% identity, 90% coverage) of both flanks on the same lycPyr6 scaffold. The coordinates of the draft genome gaps projected onto lycPyr6 were then intersected with the RepeatMasker annotation using bedtools intersect. Draft genome gaps containing only one type of repeat on lycPyr6 were classified according to the type of repeat. In case the draft genome gaps corresponded to a region containing more than one type of repeat, the gaps were classified as ‘complex’. Finally, in case that the draft genome gaps could not be mapped unambiguously (e.g., no homology, only one flank aligned or the two flanking regions mapped to different scaffolds) or mapped to gaps on lycPyr6, they were classified as ‘not scorable gaps’ (**Figure 4a**)

We also compared how many repeats were assembled in the draft assemblies compared to lycPyr6 (**Figure 4b**) by calculating the proportion of repeat base pairs present in the draft assemblies relative to the total bp in lycPyr6. This was done for each major repeat group using the RepeatMasker table (.tbl) files; more details in **Supplementary Table S10**.

## Acknowledgements

We would like to thank Max Käller, Phil Ewels, Remi-André Olsen, Joel Gruselius, Fanny Taborsak-Lines (SciLifeLab Stockholm) for generating 10X data; Olga Vinnere-Pettersson and Ida Höijer (SciLifeLab Uppsala) for generating PacBio data; Mai-Britt Mosbech and Anna Petri (SciLifeLab Uppsala) for doing the agarose gel plug extraction; Verena Kutschera at the National Bioinformatics Infrastructure Sweden at SciLifeLab (Stockholm) for bioinformatics advice; Muhammad Bilal for the help with the repeat library; Matthias Weissensteiner for the help with sample transfer, comments on the manuscript and helpful discussions; Douglas Scofield for help with the gap-filling analysis; Diem Nguyen, Marco Ricci, James Galbraith, Julie Blommaert, Ivar Westerberg, Jesper Boman for their comments on the manuscript; Octavio Gimenez-Palacio, David Adelson, James Galbraith, Hanna Johannesson, Jesper Boman, and Boel Olsson for helpful discussions. This research was supported by grants from the Swedish Research Council Formas (2017-01597 to AS), the Swedish Research Council Vetenskapsrådet (2016-05139 to AS, and 621-2014-5113 to MI), and the SciLifeLab Swedish Biodiversity Program (2015-R14 to AS). The Swedish Biodiversity Program has been made available by support from the Knut and Alice Wallenberg Foundation. A.S. acknowledges funding from the Knut and Alice Wallenberg Foundation via Hans Ellegren. The authors acknowledge support from the National Genomics Infrastructure (NGI)/Uppsala Genome Center. The work performed at NGI / Uppsala Genome Center has been funded by RFI/VR and Science for Life Laboratory, Sweden. The authors further acknowledge support from the National Genomics Infrastructure in Stockholm funded by Science for Life Laboratory, the Knut and Alice Wallenberg Foundation and the Swedish Research Council. Some of the computations were performed on resources provided by the Swedish National Infrastructure for Computing (SNIC) through Uppsala Multidisciplinary Center for Advanced Computational Science (UPPMAX). KAJ is most grateful for the financial support received from the Villum Foundation (Young Investigator Programme, project no. 15560), and from the Carlsberg Foundation (Distinguished Associate Professor Fellowship, project no. CF17-0248). We thank the State Ministry of Research and Technology (RISTEK); the Ministry of Forestry, Republic of Indonesia; the Research Center for Biology, Indonesian Institute of Sciences (RCB-LIPI); and the Bogor Zoological Museum (Tri Haryoko) for providing permits to carry out fieldwork in Indonesia and to export select samples. KAJ acknowledges a National Geographic Research and Exploration Grant (8853-10), the Dybron Hoffs Foundation and the Corrit Foundation for financial support for fieldwork in Indonesia.

## Data Access

### Disclosure declaration

Shawn Sullivan and Ivan Liachko are employed at Phase Genomics.

